# Miniaturised structured illumination microscopy using two 3-axis MEMS micromirrors

**DOI:** 10.1101/2022.09.12.507543

**Authors:** Peter Tinning, Mark Donnachie, Jay Christopher, Deepak Uttamchandani, Ralf Bauer

## Abstract

We present the development and performance characterisation of a novel structured illumination microscope (SIM) in which the grating pattern is generated using two optical beams controlled via 2 micro-electro-mechanical system (MEMS) three-axis scanning micromirrors. The implementation of MEMS micromirrors to accurately and repeatably control angular, radial and phase positioning delivers flexible control of the fluorescence excitation illumination, with achromatic beam delivery through the same optical path, reduced spatial footprint and cost-efficient integration being further benefits. Our SIM architecture enables the direct implementation of multi-colour imaging in a compact and adaptable package. The two-dimensional SIM system approach is enabled by a pair of 2 mm aperture electrostatically actuated three-axis micromirrors having static angular tilt motion along the x- and y-axes and static piston motion along the z-axis. This allows precise angular, radial and phase positioning of two optical beams, generating a fully controllable spatial interference pattern at the focal plane by adjusting the positions of the beam in the back-aperture of a microscope objective. This MEMS-SIM system was applied to fluorescent bead samples and cell specimens, and was able to obtain a variable lateral resolution improvement between 1.3 and 1.8 times the diffraction limited resolution.

## Introduction

Widefield fluorescence microscopy is an integral and fundamental tool for the life sciences, giving the user the ability to study live or fixed subcellular structures or organisms non-invasively with a high degree of specificity, contrast and temporal resolution [1–3]. One of the limiting factors of widefield fluorescence microscopy is that the spatial resolutions achievable are limited by the diffractive nature of light, typically limiting one to resolve objects no smaller than 250 nm laterally and 600 nm axially [3,4].

To image structures with dimensions less than the optical diffraction limit, an extensive range of super-resolution microscopy approaches have been developed over the past 25 years. These can be generally categorized as either localization-based or structured-based super-resolution microscopy. One of the main structured--based approaches to achieve super-resolution in live cell specimens is structured illumination microscopy (SIM). The SIM technique is based on fluorescence microscopy widefield spatial resolution doubling in 2D or 3D through the application of optical interference, creating a spatially modulated excitation light fringe pattern in the sample [5,6]. This can be achieved in live cell imaging experiments, whilst retaining the typical widefield advantages of low light doses and high temporal resolution capabilities, while avoiding the stringent requirements on sample sparsity, morphology or usage of special labels which are characteristic of other techniques [7,8].

The SIM principle was first reported by Gustafsson [5] and Heintzmann [9] and has since seen many applications in biological research [10–12]. By projection of a fine sinusoidal grating pattern onto a fluorescent specimen, the superimposition of the illumination pattern on the specimen results in the generation of Moiré fringes which contain high frequency spatial information that is outside the optical transfer function of the imaging objective.

In order to extract the new, higher resolution image information, images with different grating phase must be acquired to uniformly illuminate the full imaging area. Additionally, as the resolution improvement is only observed in the direction normal to the illumination grating, the grating must be rotated at least a further two times by 60 degrees to generate a more uniform in-plane resolution enhancement (smaller angles and/or more phase steps can be used, though this results in greater number of required rotations and phase positions). A raw 2D SIM dataset consists of a minimum of 9 images. After computational reconstruction the result is a single image which has a near isotropic doubling of the lateral widefield resolution [5,7,13–16].

The generation of SIM pattern gratings has generally been achieved using one of two methods. The first uses a fixed diffraction grating, placed in the excitation beam path, which is mechanically manipulated to utilize the 1st and 0th order diffracted beams for generating interference [5,6]. However, the mechanical manipulation process generates a limit on the achievable temporal resolutions [13,15] from the system. The second method uses spatial light modulators (SLM) or digital micromirror devices (DMD) to create tailored spatial excitation patterns in the back-aperture of a microscope objective, thereby generating the required interference patterns in the sample [8,15,17–19].

The DMD approach has been an attractive option due to the relatively low cost of the hardware, fast response times of the micromirror arrays and the absence of a requirement to display the inverse of the desired pattern in between exposure steps, as is the case with SLMs [19,20]. The DMD approach to carrying out SIM however has some limitations, For instance, the “on” and “off” positions of the individual mirror pixels create a sawtooth-like surface which requires the DMD to be treated as a blazed grating. To compensate for this blazed grating effect, the illumination must arrive on the DMD at a wavelength-dependent incident angle to achieve the greatest excitation grating efficiency by equalizing the intensity between the diffraction orders which, if not carried out will be detrimental to the grating contrast obtained [18,19,21]. Additionally, the ensuing requirement to build the microscope around this incident wavelength defined blazed angle means that multi-colour imaging is difficult to implement. However, there has been recent work showing that this is possible using computationally determined grating patterns and varying the incident angles of different laser wavelength onto the DMD [22] or thermally tuning the output wavelength of a diode laser so as to share the same blaze angle as a second laser line [23]. The other limitation of a DMD or indeed any grating-based SIM instrument is that the optical path defines the separation between the ± 1st order beams, which must match or be smaller than the back-aperture of the imaging objective lens. This defined beam separation means that if another objective lens with a different back-aperture size was needed to be used the optical beam paths would have to be changed

In part to address this, approaches that use individual control of the two in the sample plane interfering beams have been demonstrated recently using galvanometric or piezoelectric scan mirrors and piezoelectric stages for control of the grating orientation and phase, respectively [24–26]. Either two sets of two-axis galvanometric scanners are employed to position the two interference beams, or a single scanning mirror and retroreflecting prism are utilized, requiring precise alignment of multiple elements to space the beams appropriately. The independent control of the beams and use of achromatic elements allows direct multi-colour imaging.

We present an alternative method to achieve the generation of an excitation grating pattern in a fluorescent specimen through a pair of three-axis single-crystal silicon micro-electro-mechanical system (MEMS) scanning micromirrors. MEMS have been used extensively in confocal endoscopy [27–30] and are becoming popular in further optical microscopy approaches. As such they have been used to reduce the footprint and cost of a light sheet microscope [31] and also for generating homogenous widefield illumination in order to correct for intensity-based non-uniformity of photo-switching events in single molecule localization microscopy [32].

A pair of commercially available electrostatically actuated three-axis MEMS micromirrors are used to generate a controllable spatial interference pattern at the sample plane of a custom SIM system and manipulate the phase, angular orientation and pitch of the pattern for obtaining SIM images of both calibration and biological specimens. We describe the construction of the MEMS-SIM microscope and analyse its performance and capabilities.

## Materials and Methods

### MEMS structured illumination microscope design and construction

The MEMS-SIM microscope schematic is shown in Figure 1, using two 3-axis MEMS micromirror (MEMS 1 and MEMS 2) and a knife edge prism (KEP) to generate a 2D SIM excitation pattern in the sample. A λ = 532 nm laser module (OFL17-F2, OdicForce) or a λ = 473 nm solid state laser (LSR473NL-50-PS-II, Lasever Inc) are used as the excitation source. Their beam paths are combined and passed through a linear polariser with the transmission axis aligned orthogonal to the plane shown in the schematic in Figure 1. The polarised laser input was split by a broadband 50/50 beam splitter (BSW04, Thorlabs) directing the two beams towards the two 2 mm diameter aluminium coated MEMS micromirror (A7M20.2-2000AL, Mirrorcle Inc) with an incidence angle of 22.5°. The MEMS are mounted in 3D printed adaptors on 5-axis holders (3 x DT12/M, 1 x KM05FL/M, Thorlabs). The two beams reflected by the MEMS are magnified and combined through a 4f lens configuration consisting of two f = 30 mm achromatic lenses (AC127-030-A, Thorlabs), a knife edge prism (KEP, MRAK25-P01, Thorlabs) and a f = 125 mm achromatic lens (AC254-125-A, Thorlabs). The 30 mm lenses are additionally acting as scan lenses for the two MEMS, with the KEP placed close to the focus of the two beams to allow a wide positioning and tuning potential. A f = 75 mm achromatic lens (AC254-075-A, Thorlabs) then focuses the two beams telecentrically onto the back-aperture of a 100x/1.25NA microscope objective (100x Zeiss A-Plan Oil Immersion Objective). A custom aperture has been placed at the focal point of the f = 125 mm lens to clean up any scattered light propagating through the system from overfilling of the MEMS. Between the 75 mm lens and the objective, a multiband dichroic mirror (DC/ZT375/473/532/635rpc-UF1, Chroma) reflects the two beams to create an inverted microscope geometry. The two beams undergo interference at the specimen plane, generating a sinusoidal interference grating pattern with spatial frequency and orientation dependent on the beam positions at the objective back focal plane, controlled by the 3-axis MEMS.

**Figure 1:**
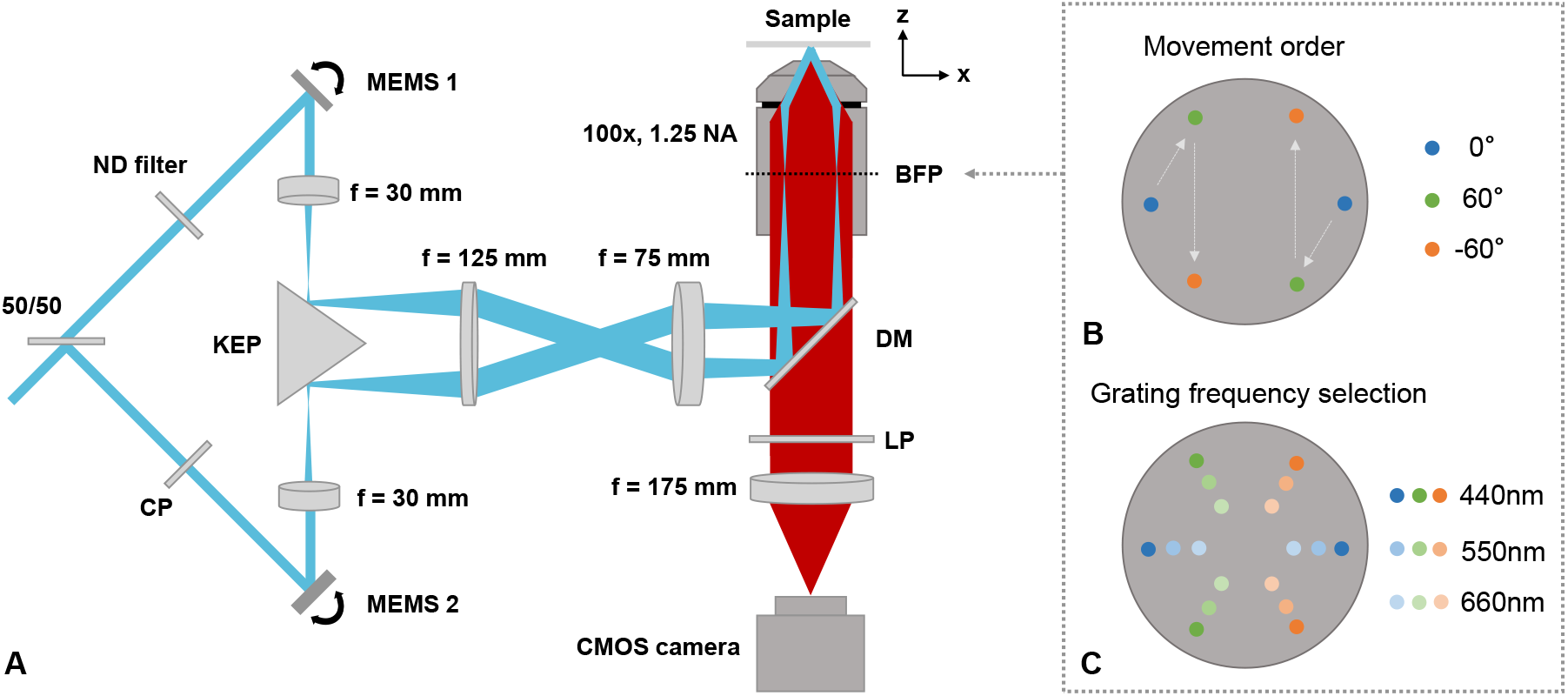
(A) Schematic of the MEMS-SIM setup using two 3-axis MEMS scanning micromirrors. All lenses are achromatic doublets. 50/50: broadband beam splitter; ND: continuous varying ND filter; CP: compensation plate; KEP: silver coated knife edge prism; DM: dichroic mirror; LP: longpass filter; BFP: back focal plane. (B) Position of the two individual controllable excitation beams focused at the back focal plane, indicating movement pattern for the 0°, 60° and −60° angles required for SIM reconstruction. (C) Back focal plane position changes to control the pitch of the excitation grating pattern, with 660 nm, 550 nm and 440 nm grating pitched used here.

The resulting fluorescence emission is collected by the objective before passing through the dichroic mirror and a multi-bandpass emission filter (69401m, Chroma) and finally being imaged onto an industrial CMOS camera (UI-3060CP-M-GL Rev.2, IDS) using a f = 175 mm achromatic tube lens (#47-644, Edmund Optics).

The specimens are placed onto a custom 3-axis sample positioning stage that consists of both 3D printed and off-the-shelf components, allowing in-plane manual positioning (2x MT1A/M, Thorlabs) and utilizing a piezo controlled focusing axis (NFL5DP20S/M, Thorlabs). The entire footprint of the system fits onto a 300 mm x 450 mm breadboard (MB3045/M, Thorlabs)

Three-axis positioning control of the MEMS is achieved through custom electronics. To allow simultaneous control of the four movement actuators of each MEMS, an 8-channel DAC is used (EVAL-AD5676, Analog Devices) in combination with two 4-channel 225 V amplifier (HV56264, Microchip Technology). The DAC is controlled using an Arduino microcontroller, which is synchronised using a custom python GUI. Using this custom electronics approach allows control of the MEMS for tip, tilt and piston movement, which is not readily possible using the commercial driver boards supplied by the MEMS manufacturer.

The overall microscope system makes use of mostly off-the shelf elements together with some custom 3D printed components and electronics, keeping the system component cost for the entire microscope including sample stage, camera, two colour laser sources and drive electronics below £9000.

### MEMS characterisation

To evaluate the quasi-static angular scan of the MEMS, a low power red laser is reflected off the mirror surface at around 10° incidence angle and the displacement of the reflected spot is measured on a screen at 60 cm distance. The angular response to DC actuation voltages is then calculated based on simple geometry. The piston and step-response movement are characterised using a microscope coupled single-point laser Doppler vibrometer (OVF512, Polytech) with vertical displacement resolution of <10 nm. The laser beam is directed to the centre of the MEMS mirror and a slow, 1 Hz stair-case voltage is applied to the MEMS, with the resulting height displacements of the micromirror measured by the vibrometer.

### Specimen preparation

All cell samples are commercially available fixed samples (FluoCells Prepared Slide #1, Invitrogen) while the 175 nm nanobead samples are prepared from solution. For these, 5 μl of 175 nm orange and green nanobeads from a PS-Speck Point Source Kit (P7220, Invitrogen) are placed on a #1.5 cover slip and air dried. Once dry, a small drop of glycerol is placed on the cover slip and a microscope slide is used to seal the sample.

### Image acquisition and processing

Raw images were acquired using a custom python GUI for MEMS control in combination with micromanager [33]for camera readout and timing. All images use 2×2 camera binning to increase the light sensitivity of the industrial CMOS camera. The number of frames to be acquired, exposure time, time delay between frames and MEMS position for all 9 SIM orientation and phase combinations is sent to the microcontroller, which will enable MEMS movement, as well as timing, control and synchronisation of the camera exposure via TTL output pulses. The dual colour exposure using the two lasers is running sequentially with a full dataset captured at a time. The acquired time series are saved in ome.tiff format, with the full camera field of view of 968×608 pixel cropped to 512×512 pixel for easier post-processing and reconstructed using Fiji [34] and the fairSIM plugin [14]. For each reconstruction a background subtraction is used followed by an approximation of the optical transfer function using the peak emission wavelength and objective numerical aperture values. Parameters for the reconstruction are estimated using a variable region to exclude from the fit (0.3 for 660nm grating period, and 0.4 for 550nm and 440nm grating period). Reconstruction is then completed using the Wiener filter option in fairSIM, with a filter parameter of 0.5 and an apodization cut-off related to the estimated resolution improvement from the parameter estimation.

## Results

### 3-axis MEMS characterisation

Before incorporation into the SIM, the two MEMS mirrors were characterized for their angular and piston movement. To enable piston movement of the commercially available mirrors, a minor modification to the bond-wires of one movement axis was undertaken, leading to a reduction of movement angle in one of the otherwise identical axes. Both angular movement response axes are shown in Figure 2A, detailing a ±10° angular range at the full Y-axis and a ±5° angular range for the reduced X-axis. The angular movement is created using a differential driving scheme between the two actuators for one rotation axis around a common offset point. Figure 2B demonstrates a simulation of the push and pull angular movement of the micromirror actuators for movement around a single axis.

**Figure 2:**
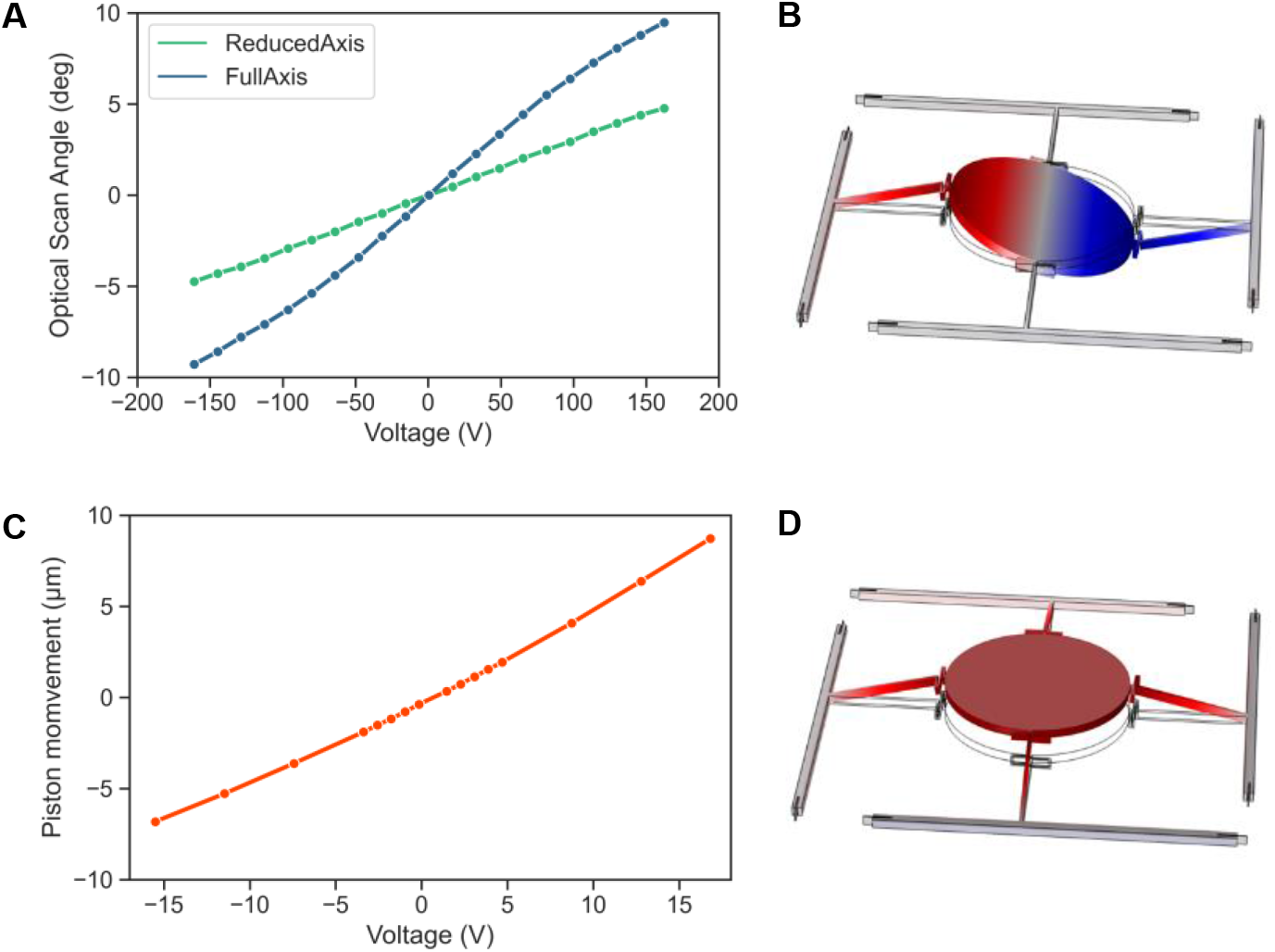
MEMS mirror characterisation. (A): Quasi-static scan angle vs DC actuation voltage for full axis (y-axis of scanner) and reduced axis (x-axis of scanner). (B): Schematic of the mirror tilt movement showing the push-pull actuation of two actuators of the same axis; (C): Piston displacement vs DC actuation voltage; (D): Schematic of the mirror piston movement showing push-push actuation.

The piston displacement of the MEMS is created through a common offset change of the actuators of the reduced angular axis only. It will translate into a phase change between the two interfering beams in the SIM microscope setup, allowing generation of the minimum 3 phase steps required for SIM reconstruction. A plot of this characterization using a reduced actuation voltage range, due to the small movements needed for the sub-wavelength phase changes required for SIM, is detailed in Figure 2C, showing a 15 μm piston movement with a 15% voltage range on the MEMS scanner actuators. Figure 2D visualises a simulation of the piston movement mode shape of the MEMS.

The step responses of the MEMS micromirrors were measured to provide information of the maximum obtainable SIM temporal resolution. The vibrometer response for two angular steps and one piston step are shown in Figure 3A to C, in each case depicting two measured steps of each response which show almost identical behaviour. The angular steps in A and B show a 10-90% step response time of less than 5 ms, but around 10 ms are required for movement oscillations of the MEMS to dampen down. In both cases a ~1.4 kHz oscillation can be seen superimposed on the step, which originates from the resonance eigenmodes of the MEMS. This oscillation can be reduced in future by using an improved higher order low-pass filter and active PID control in the MEMS actuation electronics, which will subsequently reduce the overall time to capture a single SIM frame. The piston step response (Figure 3C) has a smaller amplitude with reduced residual resonance actuation and reaches a stable position after 5 ms. The overall timing diagram for MEMS movement and synchronised camera exposure for a full SIM image, consisting of three illumination grating angle orientations with three phase steps each, is shown in Figure 3D. For the 9 images sequence pure phase steps are only present while switching phases of one grating orientation, while a return to the original phase is done with the same step as changing the grating orientation due to a single MEMS movement combining both changes. A full image with 80 ms exposure time for each individual frame used for cell imaging takes 780 ms.

**Figure 3:**
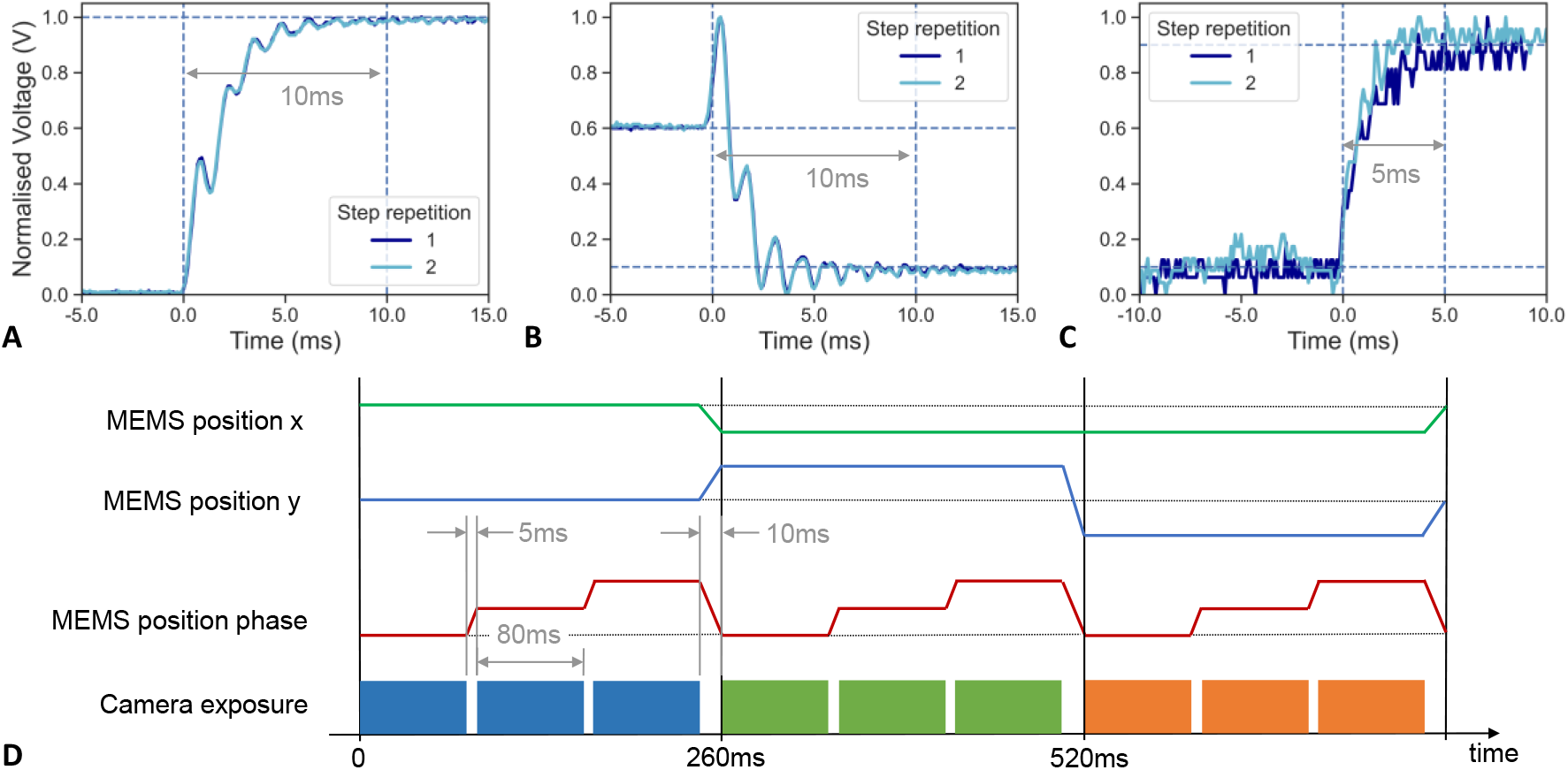
MEMS step response and MEMS-SIM timing schematic. (A) Step response for a position change from a 0° grating orientation to a 60° grating orientation. (B) Step response for a position change from 60° grating orientation to −60° grating orientation. (C) Step response for a MEMS piston movement step, changing the grating phase by 120°. (D) Timing diagram for the 3-axis MEMS movement during acquisition of the 9 raw SIM images required for reconstruction.

### Illumination grating phase shift using MEMS piston movement

To evaluate the achievable phase shift of the interference pattern generated in the sample, fluorescence images of a fixed BPAE cell slide were taken with the two MEMS positioned to create a grating with 60° orientation and grating period of 660 nm, 550 nm and 440 nm for both excitation wavelengths. The exact position was aligned using a 2D Fourier transform of the recovered fluorescence image, allowing instant access to the grating angle and period in Fourier space. Due to the variation in excitation wavelength and therefore wavelength dependent grating period, a separate position optimisation was performed for each laser. At each position a piston actuation from 0 V to 0.41 V was applied to one of the MEMS, with voltage step sizes down to 17 mV, and nine images taken at each step interval. The phase shift of the grating position between subsequent images was evaluated using a complex Fourier transform and recovering the phase of each image through an inverse tangent relationship of the Imaginary and Real part at the position of the grating response. The measured phase shift for both lasers is shown in Figure 4, with a linear response of the phase change with applied piston voltage on the MEMS visible and the phase shift for different grating periods of the same laser being in very close agreement. For the 473 nm laser a phase shift of 2/3π is achieved with 180 mV piston actuation and a 4/3π shift with 360 mV piston actuation, with a standard deviation of the measured phase step of up to 158 mrad and minimum phase step size, determined by the 16bit DAC resolution, of 35 mrad. For the 532 nm laser the equivalent shifts are achieved with 190 mV for a step of 2/3π and 380 mV for a step of 4/3π, with a standard deviation of the measured phase step of up to 70 mrad and the same DAC limited minimum phase step of 35 mrad.

**Figure 4:**
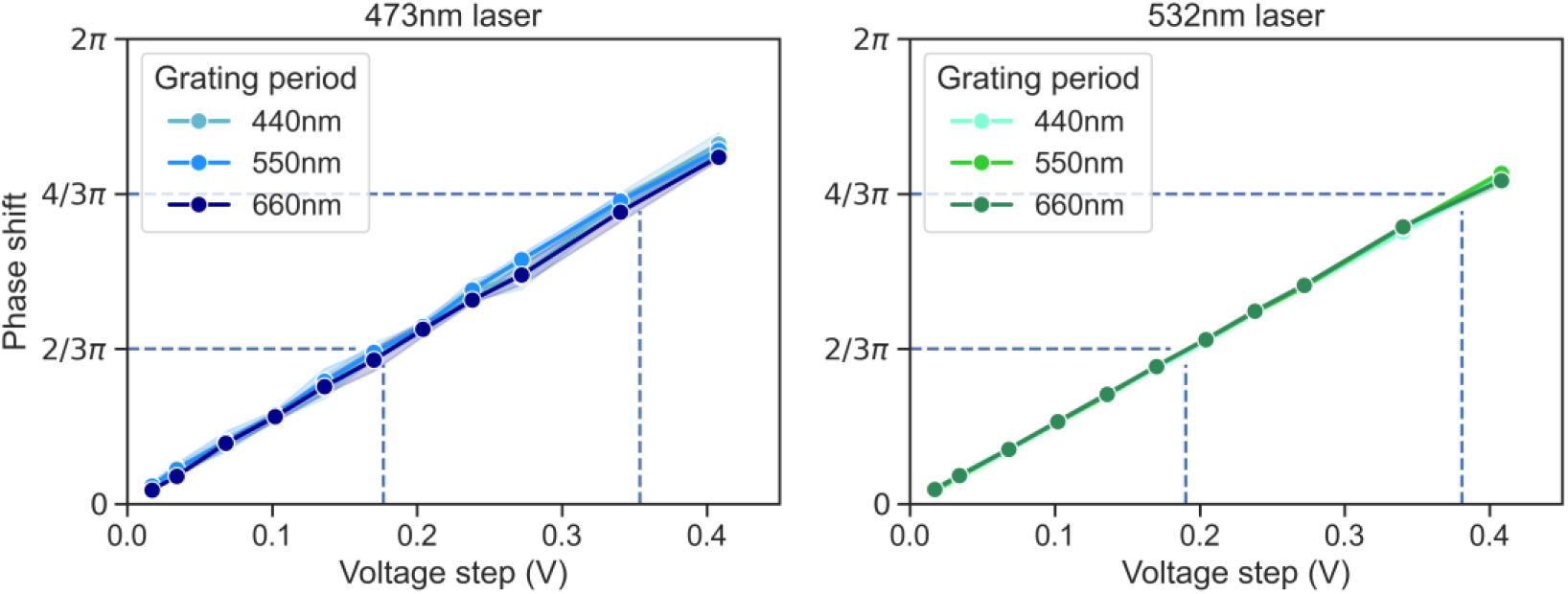
Measured phase shift of the excitation grating pattern for both 473 nm laser excitation and 532 nm laser excitation. The standard deviation of 4 measurements is shown in the banding around each line plot, with maximum phase standard deviation of 9° for the 473 nm laser and 4° for the 532 nm laser.

The grating contrast and grating phase steps of 2/3π and 4/3π are shown in Figure 5 using a fixed BPAE cell with 473 nm laser excitation wavelength and 80 ms exposure time. The overview of the camera cropped image in Figure 5A shows the 60° orientation of an excitation grating with 660 nm period. The zoom-in areas in Figure 5B show exemplar areas of all three grating orientations, with the related fringe intensity profile in Figure 5C highlighting the three phase steps required for reconstruction.

**Figure 5:**
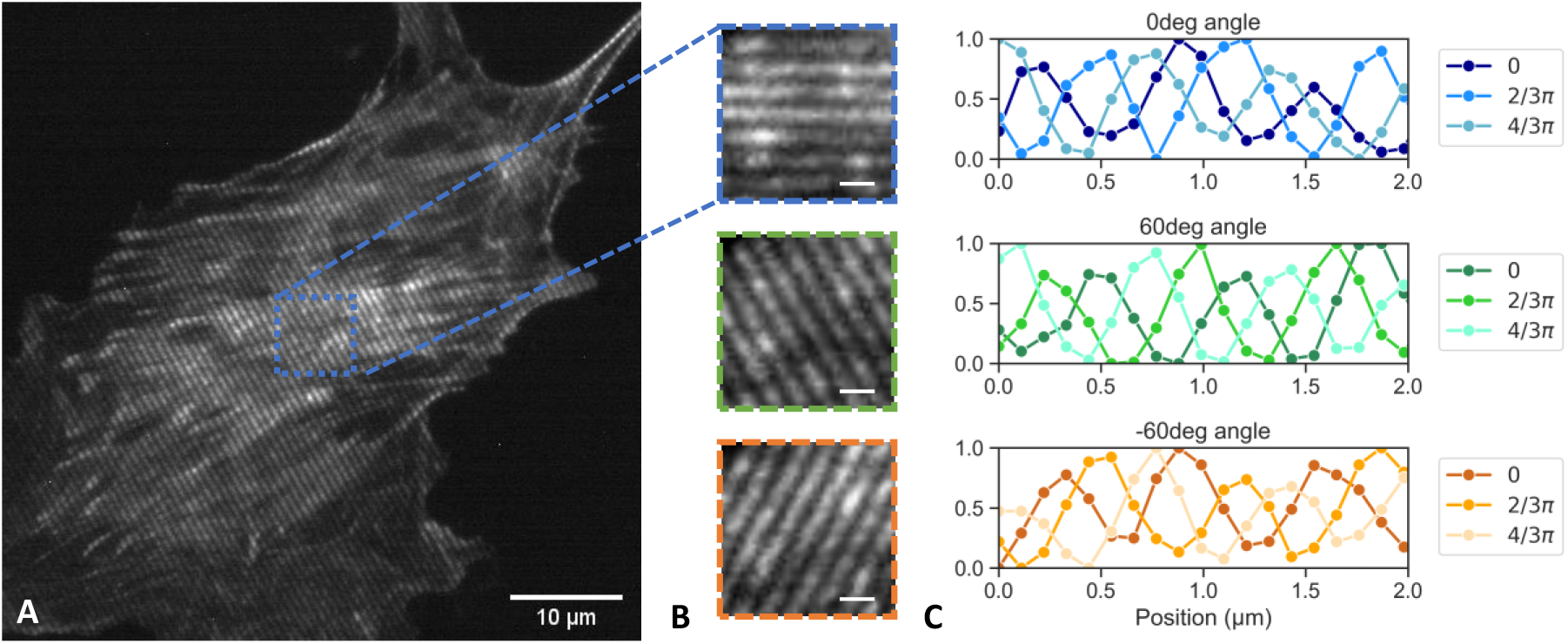
Image of 660 nm grating with 473 nm laser excitation. (A) Full field of view (B) Insets with 3 angle orientations (C) Sinusoidal intensity modulation of 3 phase steps for each grating orientation. Scale bars are 10 μm for (A) and 1 μm for (B).

### MEMS SIM imaging results

In order to quantify any lateral resolution improvement achieved using the MEMS-SIM system it was first necessary to determine the achievable widefield resolution of the custom microscope assembly. A 200 nm TetraSpeck nanosphere specimen (T14792, Invitrogen) was imaged with one of the 532 nm laser excitation beams blocked prior to the KEP using no camera binning, leading to an effective pixel size of 55 nm. Line intensity profiles were taken through 10 beads to determine the average full width at half maximum (FWHM) resolution of the system which resulted in a lateral resolution of 304 nm. Using the theoretical value of our lateral point spread function (PSF), calculated using λ/2NA where λ is approximately 550 nm and the NA of our objective is 1.25, yields a result of 220 nm which is significantly smaller than experimentally measured. However, due to the size of the beads used for imaging being of a finite size rather than significantly smaller than the diffraction limit, a correction factor must be applied to our experimentally determined PSF [35][Once this correction has been applied a modified experimental PSF of 230 nm is obtained which is in good agreement with the theoretical value.

To determine the overall resolution enhancement achievable with the MEMS-SIM system, multiple single colour fluorescence nanobeads were fixed on cover slips and their FWHM determined. Green 175 nm diameter nanobeads with excitation maximum at 505 nm and emission at 515 nm were used with the 473 nm laser, while orange nanobeads with identical diameter and excitation maximum at 540 nm and emission at 560 nm were used with the 532 nm laser. Sequences of nine images with varying angle and phase were taken with exposure times of 20 ms and laser power at the sample of 0.3 mW for the 473 nm laser and 0.2 mW for the 532 nm laser. SIM images are reconstructed using fairSIM, with figure 6 showing the comparison of pseudo-widefield (WF) images, generated from summation of all nine grating images, and reconstructed bead images for each laser with 660 nm excitation grating and 440 nm excitation grating. Bead cross sections of two closely spaced 175 nm beads are shown in figure 6C and F for all three grating periods used and both lasers. The cross-section for WF shows that the beads cannot be resolved into individual entities, while the improvement in resolution for the different SIM reconstructions with varying grating period is clearly visible. To estimate the achievable resolution, ten beads distributed over the camera field of view were measured for WF and all grating spacings. FWHM values of a Gaussian fit through the cross-section data lead to a resolution with the 473 nm laser excitation of 289±7 nm for WF and down to 176±9 nm for the reconstruction using the 440 nm grating. With 532nm laser excitation the measured resolution is at 305±8 nm for WF and down to 173±5 nm for the reconstruction using the corresponding 440 nm grating. The corresponding resolution enhancement factors were calculated as 1.6 and 1.8 respectively. An overview of the resolution values for all three grating periods is summarised in Table 1.

**Figure 6:**
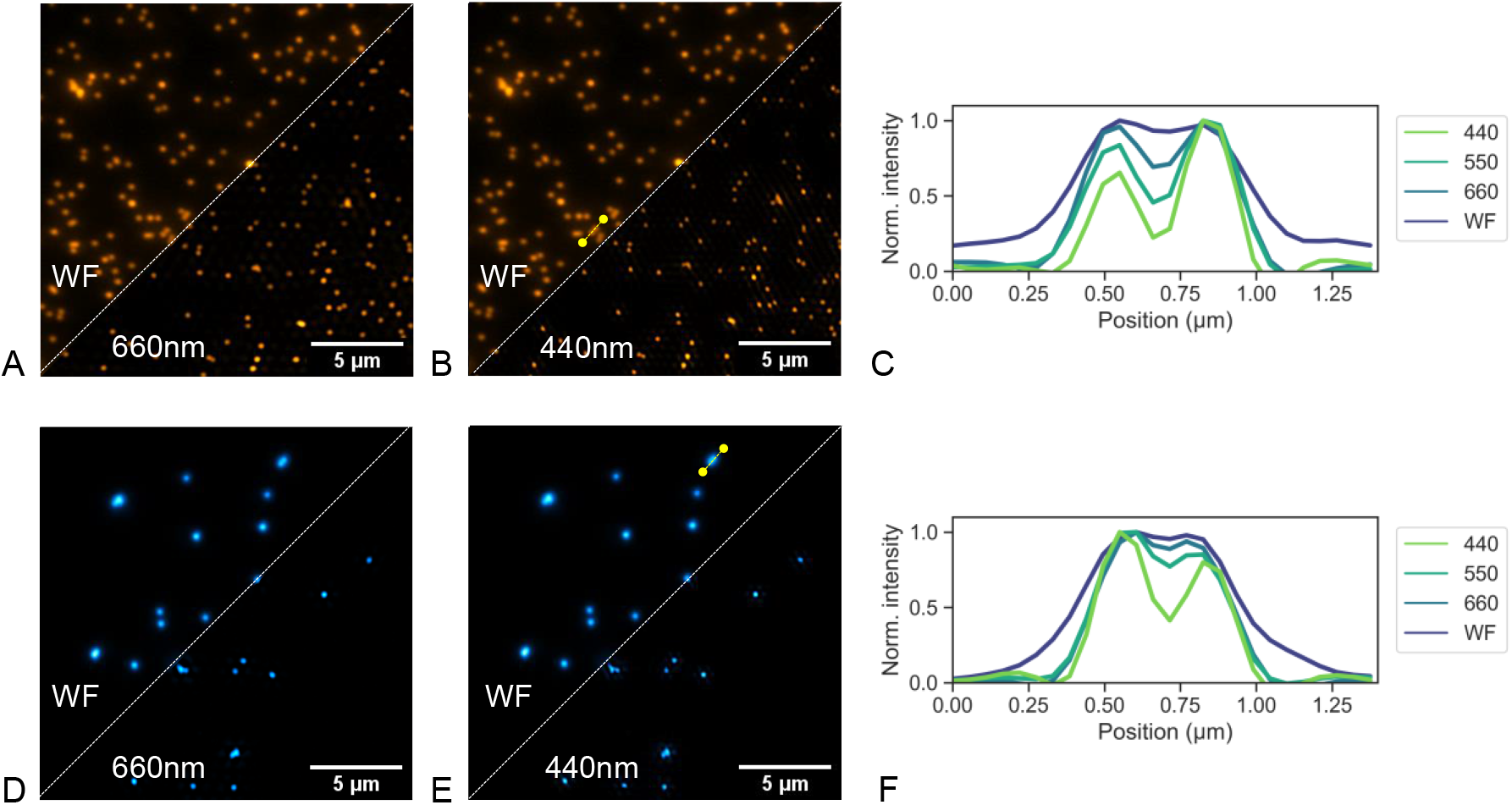
Images of sub-resolution nanobeads for both excitation wavelengths. The Images show the summed widefield images compared to SIM reconstruction images for both moderate resolution enhancement (660 nm grating period) as well as the maximum here used resolution enhancement (330 nm grating period). (A) 532 nm laser excitation with 660 nm grating SIM reconstruction, (B) 532 nm laser excitation with 440 nm grating SIM reconstruction, (C) Cross-section comparison of two neighbouring orange 175 nm beads ranging from widefield illumination to 440 nm grating SIM reconstruction. (D) 473 nm laser excitation with 660 nm grating SIM reconstruction, (E) 473 nm laser excitation with 440 nm grating SIM reconstruction, (F) Cross-section comparison of two neighbouring green 175 nm beads ranging from widefield illumination to 440 nm grating SIM reconstruction.

**Table 1:** FWHM measurement of fluorescent beads for varying excitation grating spacing.

| Grating period | Widefield | 660nm | 550nm | 440nm |
| --- | --- | --- | --- | --- |
| Bead FWHM (nm)<br>(473nm laser / 532nm laser) | 289±7 / 305±8 | 219±7 / 232±6 | 205±5 / 206±6 | 176±9 / 173±5 |
| Resolution enhancement<br>(473nm laser / 532nm laser) | 1 / 1 | 1.3 / 1.3 | 1.4 / 1.5 | 1.6 / 1.8 |

The imaging performance of fixed cell samples using the MEMS-SIM system was evaluated using a commercial fixed BPAE cell slide with actin stained with Alexa Fluor 488 Phalloidin for the 473 nm excitation laser and mitochondria stained with MitoTracker Red CMXRos for the 532 nm excitation laser. The dual colour images were captured sequentially, with laser powers of 0.3 mW and 0.2 mW respectively and exposure times of 80 ms. Images were reconstructed using fairSIM with the parameters shown in the methods section. Figure 7A shows the actin network of an exemplar BPAE cell with pseudo widefield image summation as well as SIM reconstructions based on excitation grating periods of 660 nm and 440 nm. Due to low signal to noise levels originating from the industrial CMOS camera, some artefacts are present. Overall, a clear resolution and contrast enhancement is visible through SIM processing. Similarly, figure 7B shows the mitochondria of the same cell, comparing again WF illumination with two grating periods and showing a clear improvement of resolution and contrast. The combined dual colour image is shown in figure 7C, comparing WF with 440 nm grating period imaging, with a cross-section for each of the colour channels highlighting the resolution enhancement between widefield and the SIM reconstruction using a 440 nm grating period.

**Figure 7:**
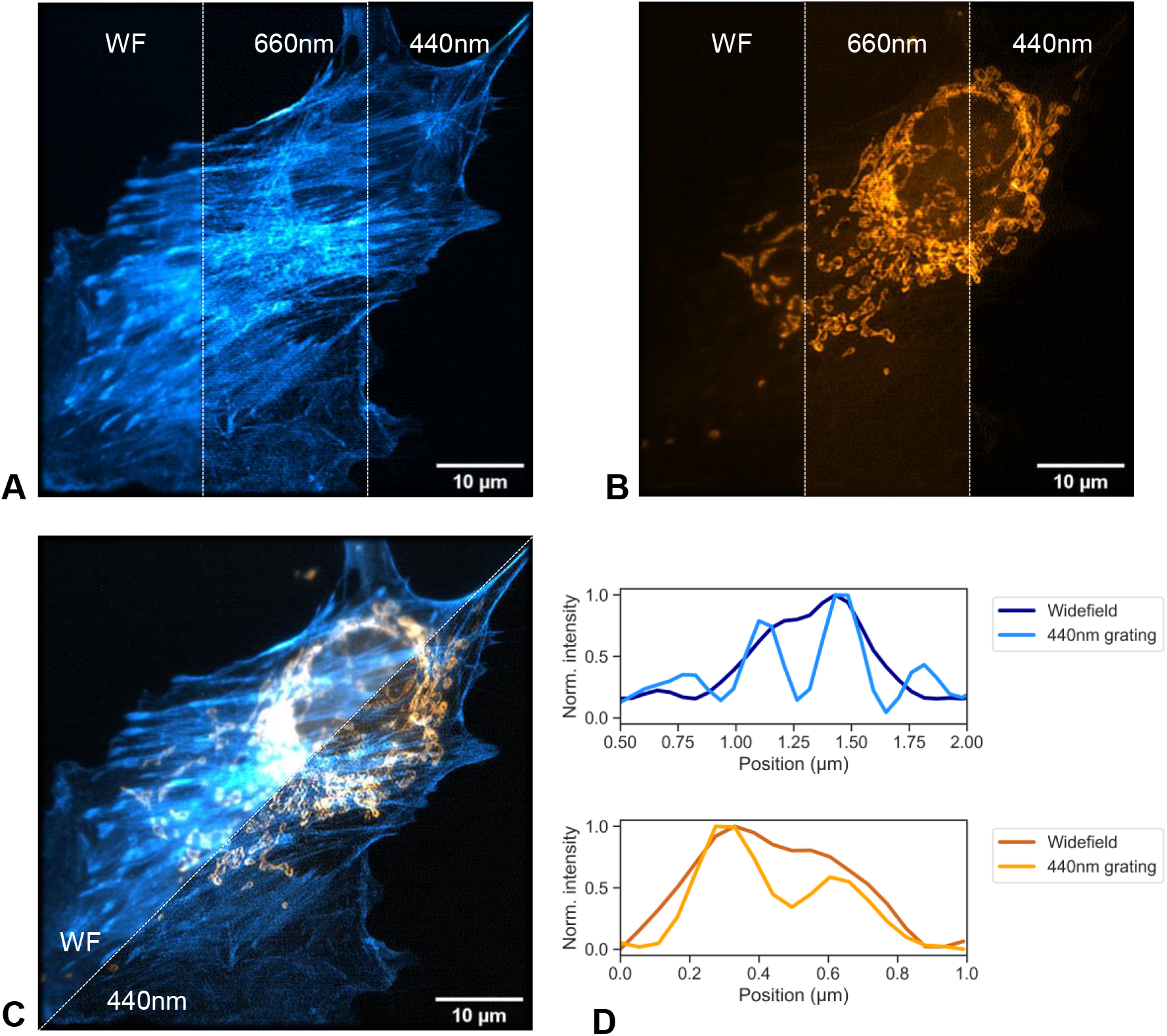
Images of fixed BPAE cells with mitochondria labelled with MitoTracke Red CMXRos and actin labelled with Alexa Fluor 488 phalloidin. Widefield images are summed result of one grating orientation. (A) Comparison of 473 nm laser excitation for widefield and fairSIM reconstructions using 660 nm and 440 nm grating period. (B) Comparison of 532 nm laser excitation for widefield and fair SIM reconstruction using 660 nm and 440 nm grating period. (C) Dual colour view of the same cell and (D) exemplary cross-sections for each colour channel, highlighting resolution improvement between widefield and 440nm grating SIM reconstruction.

## Discussion

The MEMS-SIM setup using two independent three-axis micromirrors for control of illumination grating orientation, period and phase instead of a DMD, SLM, physical grating or galvanometric mirrors reaches resolution enhancements in line with the other technologies. To balance setup simplicity and reduce background signals, no full theoretical resolution enhancement by a factor of two was aimed for, which should be reachable by including polarisation control measures in the form of a pizza polariser [20] to maintain grating contrast at highest numerical apertures. The beam positioning in the back focal plane of the 1.25NA objective used only 1/3 of the available MEMS angular displacement, as such moves to total internal reflection illumination is directly possible. With the continuous positioning control of the excitation beams in the back focal plane, swapping objectives during operation and direct integration of any high numerical aperture objective without change in any of the other parts of the physical setup is possible and an advantage of the dual micromirror approach. This is in contrast to SLM/DMD implementations where pinhole spatial filters to remove diffraction orders are used and set the physical position of the interference beams together with the beam forming optics before the objective.

Due to the non-diffractive nature of our two independently controllable beams for SIM generation, the MEMS-SIM module proves additionally to be more photon efficient than DMD or SLM based SIM systems. This can be mostly attributed to the diffraction of the excitation light by the DMD, as only 8% of this is retained between the ± 1st order beams [19]. This is further reduced by passing through the “pizza polarizer”, reducing the power by a further factor of approximately 50% [36] Not accounting for any additional optical losses through the imaging system means that optimistically 4 % of the original excitation source will propagate to the specimen. The MEMS-SIM system retains approximately 24 % of the input excitation power at the specimen plane, with losses mostly attributed to spatial filtering of any overfill of the MEMS mirror aperture in the current implementation, making it considerably more photon efficient than diffraction-based methods.

Using non-diffracting optics to generate the interference pattern creates additionally the direct ability to perform multi-colour SIM imaging as the beam paths are identical for all visible wavelength. This contrasts with DMD based SIM systems where wavelength dependent blaze conditions have to be met [21]. The main wavelength dependent requirement for the MEMS-SIM system is a broadband 50/50 beam splitter, as this will determine the power ratio in each of the interfering beams. A potential limitation of the presented setup is however related to the separation of both MEMS towards the edges of the initial beam triangle, with the physical separation of the MEMS requiring precise adjustment of the individual positions to create a path length difference that falls within the coherence length of the lasers used in the system. This can be a limitation for using low coherence laser diodes.

Currently the temporal resolution of the MEMS SIM system is limited by the MEMS step response time of up to 10 ms, as well as the signal to noise ratio on the industrial CMOS camera. The camera exposure time can be reduced to 1-20 ms if photobleaching considerations are reduced, while the MEMS step response can potentially reach <2 ms with improved filtering of the drive waveforms as well as active PID control of the angle and phase positions steps, as is the case when using galvanometric scanners. This limit is however still above theoretical step response times of DMDs, which can reach 100 μs update rates due to vacuum sealing and the discrete end stops for the binary positions of each DMD micromirror.

Aside from the imaging performance and flexibility of the presented MEMS-SIM system, there is a financial cost element to be considered. Even with open hardware approaches, such as OpenSPIM [37], costs of imaging hardware on the component level alone can easily exceed £100,000. There has been a push of late to develop low-cost imaging systems making use of industrial grade hardware and 3D-printed components, such as the UC2 [38] and open flexure systems [39]. The MEMS-SIM system was developed with this ethos in mind but aiming for metallic commercial off-the-shelf components where possible instead of fully 3D-printing to increase long-term stability. This resulted in a flexible SIM system that came in at a cost of < £9000. This is comparable to other low-cost SIM methodologies using DMDs, with the added benefit that by making use of two individual MEMS micromirrors we have full control over the angular and phase positioning without being constrained by the blaze condition of a DMD diffraction grating. This allows the MEMS-SIM approach to be used with any excitation wavelength and without objective lens back aperture constraints.

## Conclusions

In this work we have shown the design and characterisation of a structured illumination microscopy system based on the use of two 3-axis MEMS micromirror for generation of the interference gratings necessary for SIM super-resolution reconstruction. Full analogue control of the excitation grating phase, angle and frequency is shown, allowing a tailoring of the illumination pattern for optimised performance in varying samples. Characterisation of the modified MEMS mirror shows a maximum 10 ms step response time and phase control of interference gratings with down to 2° phase steps. Using commercial and 3D-printed elements to generate a MEMS-SIM system with overall component costs below £9,000 allows generation of stable and reliable SIM reconstruction with demonstrated resolution enhancement factors in its current form of up to 1.8x compared to widefield illumination. Direct multi-colour imaging has been demonstrated using microbead and fixed fluorescence cell samples.

## Acknowledgements

We acknowledge funding from the UK Engineering and Physical Sciences Research Council (grant EP/S032606/1 and studentships EP/R513349/1 and EP/T517938/1) and UK Royal Academy of Engineering (Engineering for Development Fellowship scheme RF1516/15/8).

## Data availability

All research data and materials supporting this publication can be accessed at GitHub under https://github.com/RalfBauerUoS/MEMS-SIM and at the University of Strathclyde data repository under https://doi.org/10.15129/fb83f3fa-4e7f-4049-854a-9e3325cd9779

